# Aβ-Positivity Predicts Cognitive Decline but Cognition Predicts Progression to Aβ-Positivity

**DOI:** 10.1101/523787

**Authors:** Jeremy A. Elman, Matthew S. Panizzon, Daniel E. Gustavson, Carol E. Franz, Mark E. Sanderson-Cimino, Michael J. Lyons, William S. Kremen, for the Alzheimer’s Disease Neuroimaging Initiative

**Author notes:** These authors contributed equally to the manuscript. Data used in preparation of this article were obtained from the Alzheimer’s Disease Neuroimaging Initiative (ADNI) database (adni.loni.usc.edu). As such, the investigators within the ADNI contributed to the design and implementation of ADNI and/or provided data but did not participate in analysis or writing of this report. A complete listing of ADNI investigators can be found at: http://adni.loni.usc.edu/wp-content/uploads/how_to_apply/ADNI_Acknowledgement_List.pdf. Correspondence should be addressed to Jeremy A. Elman, Ph.D., UCSD Department of Psychiatry, 9500 Gilman Drive (MC 0738), La Jolla, CA, USA, 92093.

## Abstract

**Background:** Stage 1 of the NIA-AA’s proposed Alzheimer’s disease (AD) continuum is defined as β-amyloid (Aβ) positive but cognitively normal. Identifying at-risk individuals *before* Aβ reaches pathological levels could have great benefits for early intervention. Although Aβ levels become abnormal long before severe cognitive impairments appear, increasing evidence suggests subtle cognitive changes may begin early, potentially before Aβ surpasses the threshold for abnormality. We examined whether baseline cognitive performance would predict progression from normal to abnormal levels of Aβ.

**Methods:** We examined the association of baseline cognitive composites (Preclinical Alzheimer Cognitive Composite [PACC]; ADNI memory factor score [ADNI_MEM]) with progression to Aβ-positivity in 292 non-demented, Aβ-negative Alzheimer’s Disease Neuroimaging Initiative (ADNI) participants. Additional analyses included continuous CSF biomarker levels to examine the effects of subthreshold pathology.

**Results:** Forty participants progressed to Aβ-positivity during follow-up. Poorer baseline performance on both cognitive measures was significantly associated with increased odds of progression. More abnormal levels of baseline CSF p-tau and subthreshold Aβ were associated with increased odds of progression to Aβ-positivity. Nevertheless, baseline ADNI_MEM performance predicted progression even after controlling for baseline biomarker levels and *APOE* genotype (PACC was trend level). Survival analyses were largely consistent: controlling for baseline biomarker levels, baseline PACC still significantly predicted progression time to Aβ-positivity (ADNI_MEM was trend level).

**Conclusions:** The possibility of intervening *before* Aβ reaches pathological levels is of obvious benefit. Low cost, non-invasive cognitive measures can be informative for determining who is likely to progress to Aβ-positivity, even after accounting for baseline subthreshold biomarker levels.

## INTRODUCTION

It has become clear that, because of the long prodromal period, Alzheimer’s disease (AD) treatment should begin as early as possible (1). Early intervention may be possible after identifying Aβ-positive individuals who are still cognitively normal, defined as preclinical/Stage 1 of the AD continuum proposed by the National Institute of Aging-Alzheimer’s Association (NIA-AA) research framework (2). Yet being Aβ-positive means that significant pathology is already present. It may be critically important to identify at-risk individuals *before* they develop substantial amyloid burden (i.e., at Stage 0) to improve treatment efficacy and slow progression to AD dementia. The earlier the intervention, the greater the reduction in financial and quality-of-life burden.

Examinations of AD biomarkers primarily focus on biomarkers as predictors of cognitive decline, but here our focus was on biomarker positivity as an outcome. Standard models of AD progression posit that abnormal biomarkers precede clinical symptom onset by years or even decades, and there is ample evidence to support this (3–5). However, there is also evidence that cognition may begin to demonstrate more subtle change earlier than is typically appreciated. Previous work has shown that cognition begins to show accelerated change across individuals with a range of baseline Aβ values, including those that do not meet the threshold for Aβ-positivity (6, 7). Delayed recall has been shown to demonstrate accelerating change prior to other biomarker and clinical measures (8–10). Change in amyloid is also correlated with change in cognition (11, 12). Thus, measures of Aβ accumulation, including subthreshold levels, are related to concurrent or future cognitive outcomes. However, none of these studies addressed whether baseline cognitive performance can predict progression to Aβ-positivity as an outcome. According to the NIA-AA framework staging, Aβ-positivity precedes cognitive impairment, consistent with a serial model of AD trajectories. This suggests Aβ-positivity should predict later decline in cognition, but not vice versa. Here, we tested that assumption by examining whether baseline cognition among Aβ-negative individuals could predict later progression to Aβ-positivity, even among cognitively unimpaired individuals.

Increasing evidence from autopsy studies indicates that abnormal tau appears in the brainstem during the earliest stages of AD – potentially before cortical Aβ plaque deposition – and tau is associated with poorer memory performance even in the absence of Aβ (13–16). However, individuals classified as A-/T+ are not considered to be on the AD continuum. Therefore, we also examined whether individuals with elevated tau would be more likely to progress to Aβ-positivity, which would indicate that they may eventually end up on the AD continuum, albeit with an atypical progression.

Focusing on Aβ-positivity as an outcome rather than a predictor would constitute an important step toward even earlier identification. For example, it is desirable to treat hypertension rather than waiting for the occurrence of a heart attack or stroke, but intervention aimed at preventing or delaying hypertension is even better. Similarly, being able to prevent or slow progression to Aβ-positivity is likely to be more effective in slowing AD disease progression than intervening after pathological amyloid levels have already been reached.

## METHODS

### Participants

Data used in the preparation of this article were obtained from the Alzheimer’s Disease Neuroimaging Initiative (ADNI) database (adni.loni.usc.edu). The ADNI was launched in 2003 as a public-private partnership, led by Principal Investigator Michael W. Weiner, MD. The primary goal of ADNI has been to test whether serial magnetic resonance imaging (MRI), positron emission tomography (PET), other biological markers, and clinical and neuropsychological assessment can be combined to measure the progression of MCI and early AD.

Participants from the ADNI-1, ADNI-GO, and ADNI-2 cohorts were included if they 1) had valid cognitive and cerebrospinal fluid (CSF) Aβ and phosphorylated tau (p-tau) data at baseline, 2) had at least one follow-up of amyloid data based on CSF or amyloid-positron emission tomography (PET), 3) were Aβ-negative at baseline, and 4) did not have a diagnosis of Alzheimer’s dementia at baseline (see **Table 1** for participant characteristics). In total, baseline and follow-up amyloid status were based on 585 assessments of CSF Aβ, 646 florbetapir PET scans, and 10 ^11^C-Pittsburgh Compound B (PIB) scans. Individuals were classified as Aβ-stable if they showed no evidence of abnormal amyloid at any follow-up, or as Aβ-converter if they showed evidence of abnormal Aβ at a follow-up assessment. Aβ-positivity was determined with either CSF or PET (see below). Individuals who were Aβ-positive at multiple assessments followed by a subsequent reversion to normal Aβ status on only a single timepoint were included as Aβ-converters. Individuals who were only Aβ-positive at one assessment followed by reversion to normal, i.e., Aβ-negative status, were excluded (n=9). Individuals diagnosed as MCI in ADNI (17) were included if they were Aβ-negative at baseline because our focus was to determine whether poorer cognition may precede amyloid positivity, and some of these Aβ-negative individuals with MCI may progress to Aβ-positive. Excluding these individuals would truncate the distribution of cognitive performance, which was our predictor of primary interest. A total of 292 individuals were included (252 Aβ-stable, 40 Aβ-converters). Despite being Aβ-negative, 138 (47.3%) were diagnosed with MCI at baseline.

**Table 1.** Baseline sample characteristics of Aβ-stable versus Aβ-converters. Descriptive statistics of Aβ-stable and Aβ-converter participants at baseline. Mean (SD) presented for continuous variables, count (%) presented for categorical variables. An asterisk indicates a significant (p < 0.05) difference between the two groups.

| | Aβ-stable | Aβ-converter |
| --- | --- | --- |
| n | 252 | 40 |
| Age | 71.62 (7.20) | 71.69 (6.71) |
| Gender, male | 128 (50.8%) | 25 (62.5%) |
| APOE-ε4+ status | 41 (16.3%) | 12 (30.0%) |
| MCI Diagnosis | 117 (46.4%) | 21 (52.5%) |
| Education* | 16.21 (2.56) | 17.20 (2.22) |
| Length of follow-up (years)* | 3.22 (1.59) | 4.30 (2.44) |

Procedures were approved by the Institutional Review Board of participating institutions and informed consent was obtained from all participants.

### CSF and amyloid imaging measures

CSF samples were collected and processed as previously described (18). CSF Aβ_42_ and p-tau were measured with the fully automated Elecsys immunoassay (Roche Diagnostics) by the ADNI biomarker core (University of Pennsylvania). Established cutoffs designed to maximize sensitivity in the ADNI study population were used to classify biomarker positivity [Aβ+: Aβ_42_<977 pg/mL; p-tau+: p-tau>21.8 pg/mL] (http://adni.loni.usc.edu/methods) (19).

PET Aβ was measured with the tracers PIB and ^18^F-florbetapir; PET data were processed according to previously published methods (http://adni.loni.usc.edu/methods) (20, 21). Mean standardized uptake value ratios (SUVR) were taken from a set of regions including frontal, temporal, parietal and cingulate cortices using whole cerebellum (florbetapir) or cerebellar gray matter (PIB) as a reference region. Established cutoffs to determine Aβ+ were used for PIB-PET (SUVR>1.44) and florbetapir-PET (SUVR>1.11) (20).

### Cognitive measures

We used two composite measures of baseline cognition. ADNI_MEM is based on a factor model of scores from four episodic memory tests: Rey Auditory Verbal Learning Test (RAVLT), Alzheimer’s Disease Assessment Schedule–Cognition (ADAS-Cog) word list and recognition, Mini-Mental State Examination (MMSE) word recall, and Logical Memory immediate and delayed recall (22). The Preclinical Alzheimer Cognitive Composite (PACC) (23, 24) is designed to detect amyloid-related cognitive decline and is based on Delayed Recall from the ADAS-Cog and Logical Memory, MMSE total score, and Trail Making Test, Part B time. ADNI_MEM and PACC scores were converted to z-scores and coded such that higher scores reflect *poorer* performance.

### Covariates

Age and *APOE* genotype (ε4+ vs. ε4−) were included because of their association with increased amyloid (25). Length of follow-up was included to account for decreased odds of observing an eventual progression to Aβ-positivity with shorter a period of follow-up. Education was included to account for long-standing differences in cognitive ability or cognitive reserve that might influence the relationship between amyloid and cognition. In other analyses, baseline biomarkers were included to assess the effect of AD-related pathology on progression to Aβ-positivity. P-tau status (p-tau+ vs. p-tau−) was included to account for differences in cognition due to other AD-related pathology. An additional set of models included continuously measured CSF Aβ_42_ and p-tau as covariates to determine whether subthreshold levels of pathology predict later progression to Aβ-positivity. These measures were converted to z-scores and values of CSF Aβ_42_ were reverse coded such that higher values of both measures indicated abnormality.

### Statistical analysis

We tested Aβ-stable and Aβ-converter groups for differences in the covariates using χ^2^ and t-tests. Logistic regression models were used to test whether baseline cognition in Aβ-negative individuals was associated with increased odds of future progression to Aβ-positivity. We chose this approach over a generalized linear mixed-effects (GLMM) logistic regression that includes data from all timepoints because the issue of primary interest was the odds of progressing to Aβ-positivity at any point during follow-up as opposed to the odds of being Aβ-positive at each individual timepoint (see Supplemental Material for further discussion). The first set of models separately tested the ADNI_MEM and PACC, with baseline cognitive performance on these measures as predictors and group (Aβ-stable or Aβ-converter) as the outcome. The second set of models additionally included p-tau status (p-tau+ vs. p-tau−) to assess whether lower cognitive performance was driven by abnormal levels of p-tau, the other hallmark pathology associated with AD. Although no subject met criteria for abnormal Aβ at baseline, that does not mean they were completely free of pathology. Therefore, we ran a third set of models to determine whether poorer cognition at baseline was driven by sub-threshold levels of amyloid or tau pathology. These models included levels of CSF Aβ_42_ and p-tau as continuous predictors. All models included age at baseline, *APOE* genotype (ε4+ vs. ε4−), education, and length of follow-up as covariates.

Although our primary aim was to determine whether baseline cognition was associated with increased odds of progression to Aβ-positivity at any point during follow-up rather than its association with time to progression, we sought to more directly address potential differences in follow-up time by conducting survival analyses. Cox proportional hazards models were used to test the association of baseline cognitive performance with time to (either conversion to Aβ-positive or censored at last follow-up). Two sets of models were run: the first included baseline cognitive performance as the primary predictor of interest, and the second added continuous levels of baseline CSF Aβ and p-tau. These models additionally controlled for age at baseline, *APOE* genotype, and education. Analyses were conducted with R version 3.5 (26).

## RESULTS

### Descriptive statistics

Descriptive statistics are presented in **Table 1** and **Table 2**. There were no significant differences between groups for age (*P*=0.94), gender (*P*=0.18), or proportion of individuals with MCI (*P*=0.47). Aβ-converters were more likely to be *APOE*-ε4+, but this difference did not reach significance (*P*=0.08). The Aβ-converter group had a higher average education (17.3 vs. 16.2 years; *t*=2.78, *P*=0.007). Follow-up interval was significantly longer for the Aβ-converter group (4.22 vs. 3.23 years; *t*=2.50, *P*=0.02). The mean time between baseline cognitive testing and the assessment at which Aβ-converters first demonstrated progression to Aβ-positivity was 2.8 years (interquartile range: 1.98–4.01 years). Of the 138 individuals who were Aβ-negative and had MCI at baseline, 21 (15%) progressed to Aβ-positivity. The MCI group as a whole did not have significantly different levels of baseline CSF Aβ (*P*=0.119) or p-tau (*P*=0.930) compared to cognitively normal participants. However, individuals with MCI that progressed to Aβ-positivity did have lower levels of baseline CSF Aβ (*t*=3.158, *P*=0.004) and higher levels of p-tau (*t*=2.389, *P*=0.024) compared to those with MCI that did not (see **Supplemental Table S1**).

**Table 2.** Baseline sample characteristics of cognitively normal versus mild cognitive impairment. Descriptive statistics of cognitively normal participants versus those with mild cognitive impairment at baseline. Mean (SD) presented for continuous variables, count (%) presented for categorical variables. An asterisk indicates a significant (p < 0.05) difference between the two groups.

|  | CN | MCI |
| --- | --- | --- |
| n | 154 | 138 |
| Age | 72.67 (5.97) | 70.47 (8.09) |
| Gender, male | 80 (51.9%) | 73 (52.9%) |
| APOE-ε4+ status | 25 (16.2%) | 28 (20.3%) |
| Education* | 16.50 (2.50) | 16.18 (2.57) |
| Baseline CSF Aβ* | 1488.68 (233.40) | 1443.50 (260.50) |
| Baseline CSF P-tau* | 19.47 (5.75) | 19.54 (7.86) |

### Baseline cognition predicts progression to Aβ-positivity during follow-up

In the first set of models, Aβ-converters were also more likely to be *APOE*-ε4 carriers, have more education, and longer duration of follow-up. Age was not significantly associated with progression to Aβ-positivity in either model. After accounting for covariates, individuals with poorer performance on either cognitive composite at baseline showed higher odds of progressing to Aβ-positivity at follow-up (ADNI_MEM: OR=1.66, *P*=0.013; PACC: OR=1.66, *P*=0.01). Full results of the regression models are presented in **Figure 1**.

**Figure 1.**
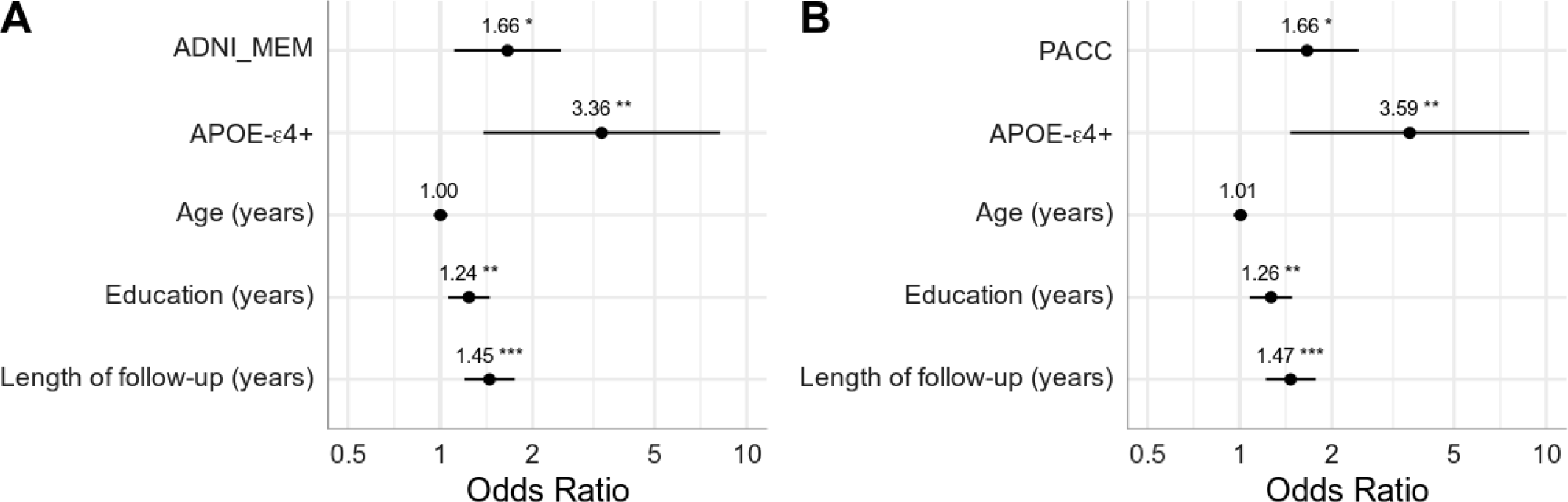
Baseline cognitive performance predicting future conversion to Aβ-positivity. Results of two logistic regression models using A) the ADNI Memory composite (ADNI_MEM) and B) the Preclinical Alzheimer Cognitive Composite (PACC). Measures are all taken from baseline and predict future progression to Aβ-positivity. Cognitive scores were converted to z-scores and reverse coded such that higher scores indicate poorer performance. Odds ratios are presented with asterisks indicating significant estimates (*p<0.05, **p<0.01, ***p<0.001). Lines represent 95% confidence intervals.

The second set of models included a dichotomous classification for baseline CSF p-tau (**Figure 2**). As in the first set of models, Aβ-converters were more likely to be an *APOE*-ε4 carrier, have more education, and longer duration of follow-up. Age and dichotomous classification of p-tau status were not significantly associated with progression to Aβ-positivity in either model. After controlling for covariates, poorer baseline performance on either cognitive composite remained significantly associated with increased odds of progressing to Aβ-positivity at follow-up (ADNI_MEM: OR=1.64, *P*=0.016; PACC: OR=1.67, *P*=0.011).

**Figure 2.**
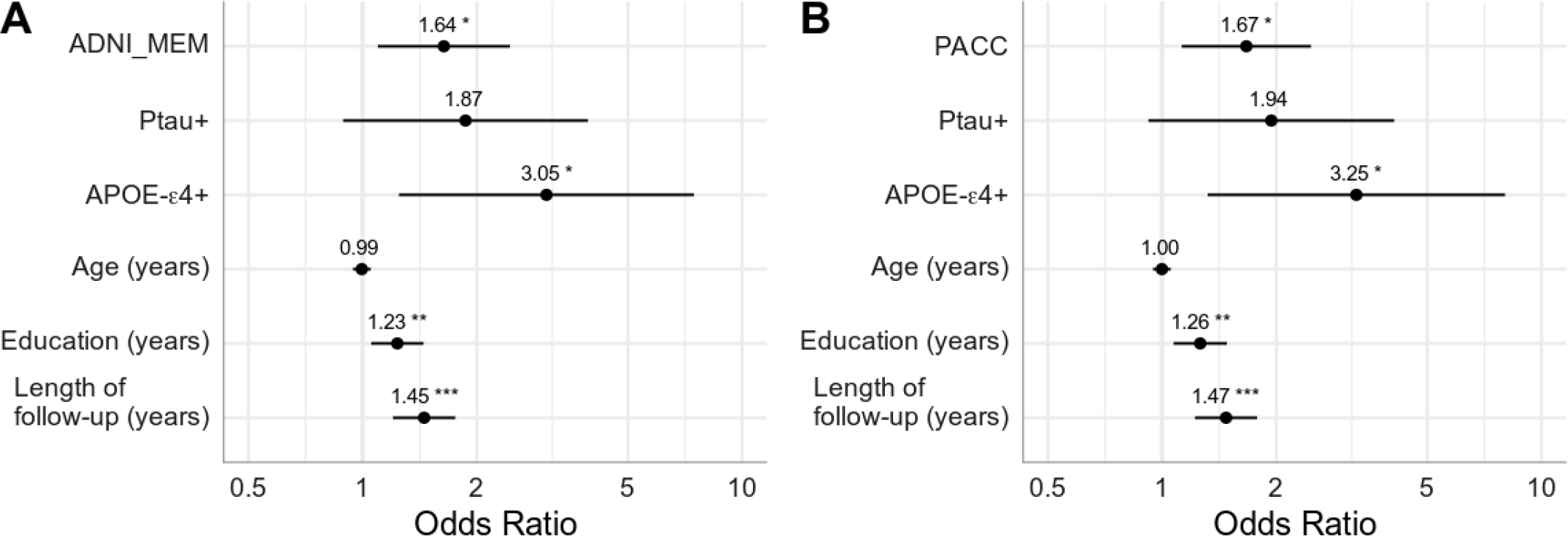
Baseline cognitive performance and p-tau+ status predicting future conversion to Aβ-positivity. Results of two logistic regression models using A) the ADNI Memory composite (ADNI_MEM) and B) the Preclinical Alzheimer Cognitive Composite (PACC). Measures are all taken from baseline and predict future progression to Aβ-positivity. Cognitive scores were converted to z-scores and reverse coded such that higher scores indicate poorer performance. P-tau-positivity is entered as a dichotomous variable. Odds ratios are presented with asterisks indicating significant estimates (*p<0.05, **p<0.01, ***p<0.001). Lines represent 95% confidence intervals.

The third set of models addressed the question of whether subthreshold levels of AD pathology could account for the effect of lower cognitive performance on progression by including continuous CSF Aβ and p-tau measures (**Figure 3**). More abnormal levels of baseline CSF Aβ and p-tau were associated with increased odds of progression to Aβ-positivity (CSF Aβ: OR=2.53 – 2.59, *P*<0.001; CSF p-tau: OR=1.51, *P*=0.03). In the case of CSF Aβ, we note that these values were all in the normal range according to standard cut-offs. After controlling for baseline biomarkers, the performance on the ADNI_MEM remained a significant predictor (OR=1.61, *P*=0.03), but the effect of the PACC was reduced to trend level (OR=1.49, *P*=0.071). Education and length of follow-up remained significant predictors of progression, whereas the effect of *APOE*-ε4 status was reduced to trend level.

**Figure 3.**
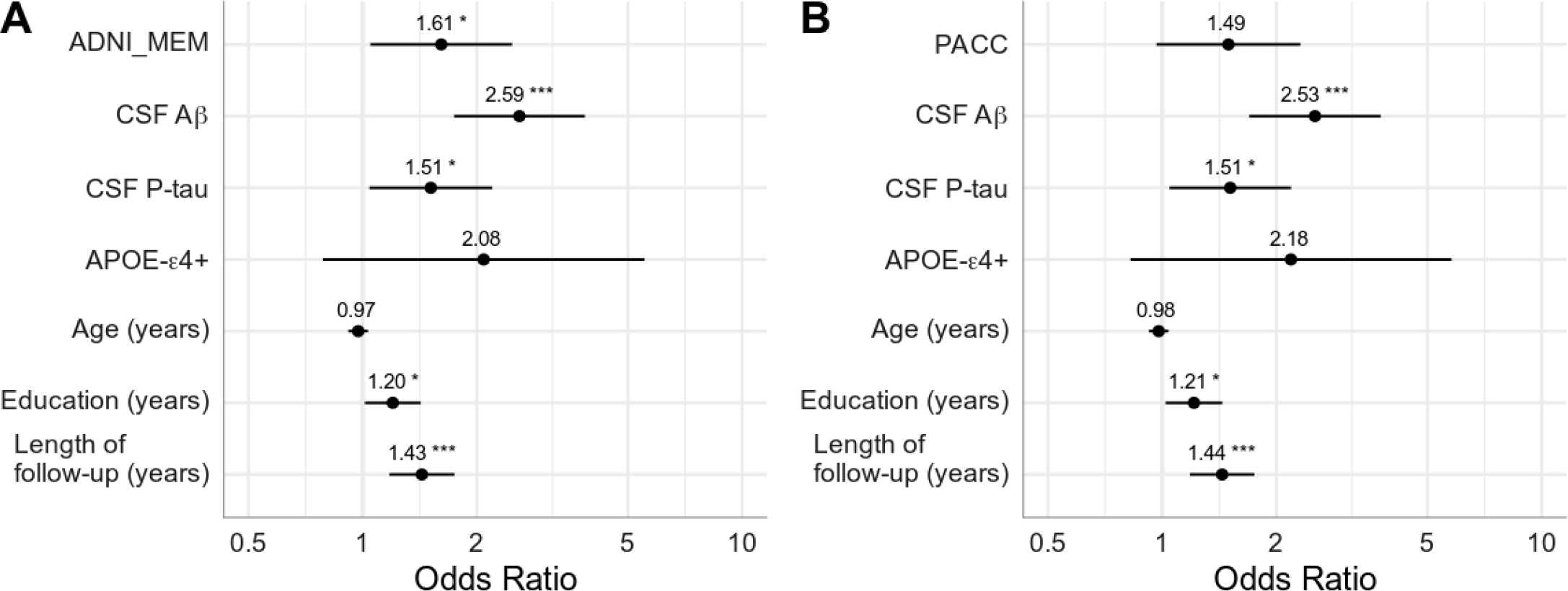
Baseline cognitive performance and continuous measures of CSF Aβ and p-tau predicting future conversion to Aβ-positivity. Results of two logistic regression models using A) the ADNI Memory composite (ADNI_MEM) and B) the Preclinical Alzheimer Cognitive Composite (PACC). Measures are all taken from baseline and predict future progression to Aβ-positivity. Cognitive scores were converted to z-scores and reverse coded such that higher scores indicate poorer performance. CSF Aβ and P-tau were entered as continuous variables. Both measures were z-scored and CSF Aβ was reverse coded such that higher values on both indicates abnormality. Odds ratios are presented with asterisks indicating significant estimates (*p<0.05, **p<0.01, ***p<0.001). Lines represent 95% confidence intervals.

To determine whether these results may be driven by the MCI participants, we conducted follow-up analyses on CN and MCI groups separately. The large drop in sample size resulted in non-significant results for most analyses, but the effects of cognition predicting progression to Aβ-positivity tended to be larger for the CN group.

### Baseline cognition predicts progression time to Aβ-positivity

The cox proportional hazard models were largely consistent with results from the logistic regression models. In models including only baseline cognitive performance and covariates, *APOE*-ε4 and higher education were associated with significantly higher risk whereas age was not. After accounting for covariates, lower cognitive performance was associated with significantly increased risk of progression to Aβ-positivity (ADNI_MEM: HR=1.48, *P*=0.024; PACC: HR=1.61, *P*=0.006). See **Figure 4** for plots of survival curves based on baseline cognitive performance and **Supplemental Figure S1** for full model results.

**Figure 4.**
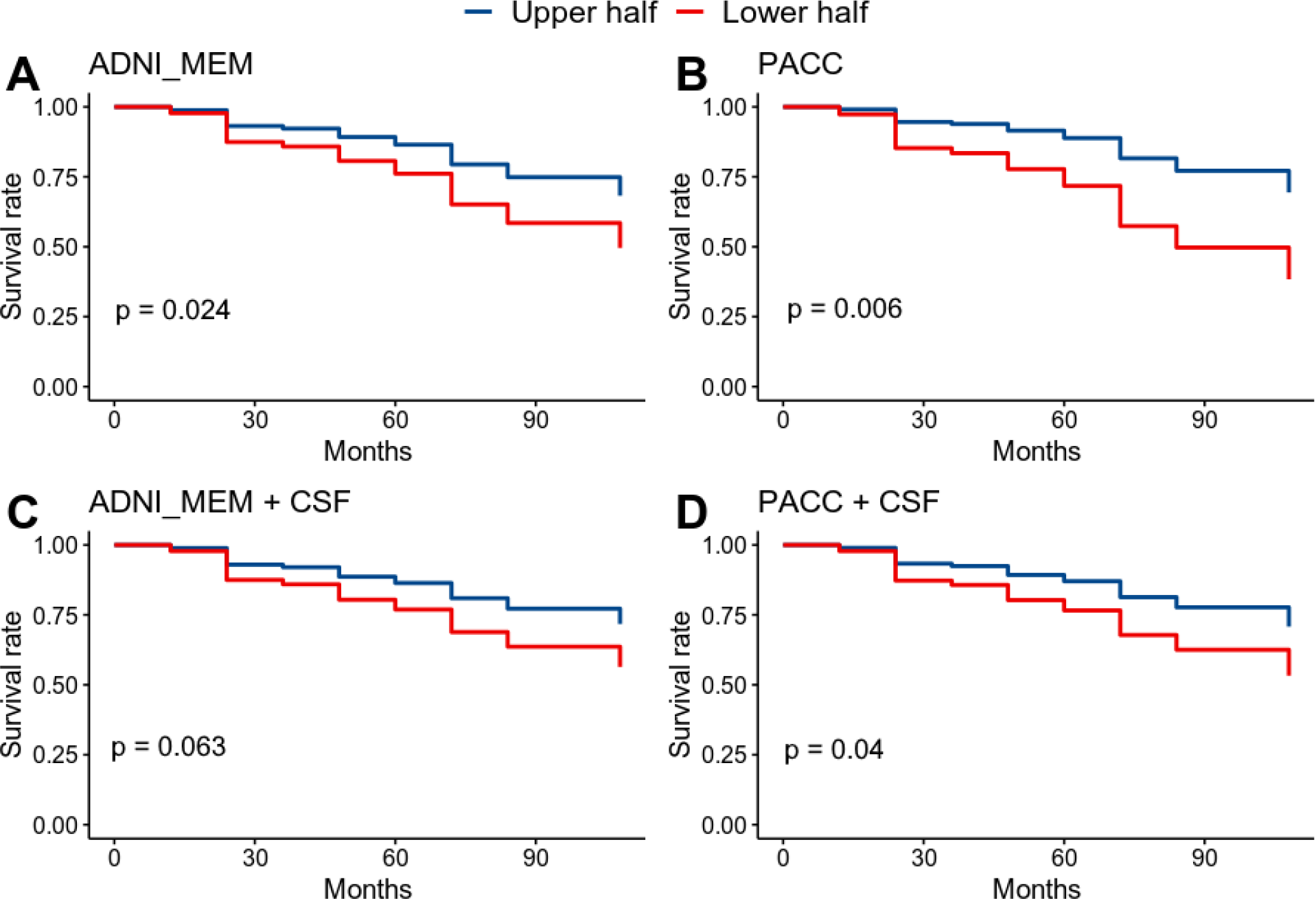
Survival estimates of progression to Aβ-positivity based on baseline cognitive performance. Cox proportional hazard models were run using continuous measures of baseline performance. For display purposes, scores were grouped based on a median split and adjusted survival curves are shown for better (upper half) and worse (lower half) performance on baseline cognitive measures. Results from 4 models are presented: A) ADNI Memory composite (ADNI_MEM) + covariates; B) the Preclinical Alzheimer Cognitive Composite (PACC) + covariates; C) ADNI_MEM + covariates + baseline CSF Aβ and p-tau; D) PACC + covariates + baseline CSF Aβ and p-tau. CSF Aβ and P-tau were entered as continuous variables. Covariates include: APOE-ε4+ status, age at baseline, and education. P-values of hazard ratios for cognitive measures are shown for each model.

Additional Cox models were conducted including baseline levels of CSF Aβ and p-tau to assess the impact of subthreshold pathology on risk of progression to Aβ-positivity. More abnormal levels of baseline CSF Aβ and p-tau were associated with increased risk of progression to Aβ-positivity (CSF Aβ: HR=2.3, *P*<0.001; CSF p-tau: OR=1.5, *P*<0.001). The PACC remained significantly associated with increased risk of progression (HR=1.45, *P*=0.04) whereas the effect of the ADNI_MEM was reduced to trend level (HR=1.41, *P*=0.063). Age was not associated with increased effects, and both *APOE*-ε4 and education were reduced to trend level. See **Figure 4** for plots of survival curves based on baseline cognitive performance and **Supplemental Figure S2** for full model results.

## DISCUSSION

### Cognitive function predicts Aβ-positivity

The ability to identify individuals at risk *before* substantial Aβ accumulation would enhance prospects for earlier intervention to slow AD progression. Here we found that in baseline Aβ-negative individuals, those with lower baseline cognitive performance were more likely to progress to Aβ-positivity at follow-up. The NIA-AA research framework represents a move toward defining AD as a biological construct (2). However, as noted by the NIA-AA workgroups on diagnostic guidelines for AD (27), behavioral markers may still hold great promise for early identification. A number of studies predicting progression from MCI to AD find that cognitive measures can predict future decline as well as or better than biomarkers (28–28). It is not surprising that cognitive measures predict future cognition, but we found that cognitive measures can also predict progression to Aβ-positivity even after accounting for baseline biomarker levels. Thus, cognition can be a useful early risk indicator.

### Impact of subthreshold Aβ

It is worth asking why cognition would predict future accumulation of AD pathology, and there may be several potential explanations. Pathological processes may already be underway, and lower cognitive function may represent a decline driven by subthreshold pathology. In a smaller (n=35) study of ADNI participants, baseline Aβ predicted later progression to Aβ-positivity but cognition did not (32). However, with the larger sample in our analysis, cognitive function was a significant predictor. Controlling for subthreshold Aβ in our analysis attenuated the effect of cognition, lending support to the idea that even low levels of Aβ are at least partially contributing to lower cognitive performance. This fits with growing evidence that subthreshold levels of Aβ are clinically relevant. In this case, it is simply that cognitive tests at this early stage are more sensitive than dichotomous classifications of biomarker abnormality at current detection thresholds. As biomarker measures become more sensitive, classification of biomarker abnormality may more consistently appear before cognitive differences.

On the other hand, cognition still predicted future progression to Aβ-positivity even after controlling for subthreshold Aβ. Therefore, cognitive performance contributes independent information, and the effect is not driven solely by individuals closer to the Aβ-positivity threshold. Cognitive testing early on is also more practical, non-invasive, and far less costly than CSF or PET biomarkers.

The relevance of subthreshold pathology also has implications for the use of dichotomous versus continuous biomarker measures. The A/T/(N) framework classifies individuals based on dichotomous biomarker measures. However, the framework authors do raise the possibility that different thresholds may be required depending on the research context (2). Some have argued that making Aβ thresholds less conservative may improve sensitivity without a substantial sacrifice of specificity (33). Our results suggest that analysis of continuous measures should be conducted when possible because continuous and binary A/T/(N) measures may lead to inconsistent inferences. An alternative approach is to examine accumulation of Aβ over time. Several studies have examined individuals who do not meet the criteria for abnormal Aβ but do demonstrate evidence of change in Aβ (11, 12, 34–36). These studies find that a change in levels of Aβ is correlated with concurrent cognitive decline. This decline in cognitive performance is commonly assumed to result from Aβ accumulation. Here we shifted the focus earlier in time and found that baseline cognition itself can predict later Aβ accumulation.

### Non-AD-related processes and the ordering of AD-related changes

An alternative explanation for cognition predicting Aβ-positivity is that lower cognitive function at baseline may be the result of a non-AD-related process. Individuals who progress to MCI while being Aβ-negative exhibit different biomarkers and cognitive profiles and tend to be on a non-AD trajectory (37). Consistent with this, the total MCI group in our analysis did not differ from the cognitively normal group on baseline Aβ or p-tau, perhaps suggesting a non-AD etiology for cognitive impairment. However, the significant association between baseline cognition and later Aβ-positivity suggests that such processes are still somehow a risk factor for AD. Indeed, 15% of Aβ-negative MCI participants in the present study did progress to Aβ-positivity, at which point this subset would be classified as Stage 3 in the AD continuum. This 15% had more abnormal levels of baseline Aβ (although still subthreshold) and p-tau compared to MCI participants that did not progress, suggesting that AD pathology may at least partially contribute to their cognitive impairment. Some individuals may be more sensitive to the effects of Aβ such that even subthreshold levels result in cognitive impairment.

It is, of course, possible to have mixed etiology driving impairment, regardless of whether it appears before or after an individual surpasses the threshold for Aβ-positivity. Although the A/T/(N) framework is agnostic to the sequence of AD-related changes (38), these Aβ-negative (A-) MCI cases would not be considered to be on the AD continuum. As such, there may be a tendency to assume that when it precedes Aβ-positivity, cognitive impairment must have a non-AD etiology. However, as pointed out in the NIA-AA framework, it is also uncertain that cognitive impairment arising after Aβ-positivity is solely due to AD pathology (2). Indeed, it is well known that there can be significant AD pathology without cognitive impairment (39–39). Therefore, although the proposed NIA-AA research framework staging captures the typical progression, it will be beneficial to maintain a degree of flexibility to account for individuals who may progress through these stages in a non-typical trajectory.

Tau-PET studies find that tau is confined to the medial temporal lobe and only spreads to the rest of the isocortex once Aβ is present (42–45). However, some have suggested that tau and Aβ develop independently, which may give rise to variable ordering in their progression (14, 15, 46). These different findings may raise questions about serial models of AD biomarker trajectories, i.e., that Aβ always precedes tau. We found that continuous – but not dichotomous– levels of CSF p-tau were associated with significantly higher odds of progression to Aβ-positivity. Thus, some individuals with elevated tau and subthreshold Aβ do develop typical AD-like profiles. Being at heightened risk of entering the AD continuum, they would be worth monitoring more closely.

### Long-standing individual differences

Another explanation for why cognition predicts Aβ-positivity is that lower baseline cognition might reflect long-standing individual differences. Lower cognitive function may reflect less efficient neural processing, which would in turn require higher activity. It has been proposed that elevated synaptic activity across the lifespan could result in increased release and aggregation of Aβ (47). Individuals with less efficient processing (indexed by lower cognitive function) may therefore be at greater risk of accumulating Aβ.

### Impact of educational attainment

In an unexpected finding, higher education was associated with increased odds of progression to Aβ-positivity. We propose two potential explanations. First, individuals with lower education may be at greater risk of becoming Aβ-positive prior to their baseline visit, and thus would not have been included in our analysis. Those with lower education who remained Aβ-negative up to the age of their baseline visit may be more resistant to Aβ deposition, and thus less likely to progress in the future. Second, the seemingly paradoxical education finding might be, in part, a function of ADNI ascertainment. Average education was 16+ years, yet only about 10% of this age cohort in the U.S. attained a 4-year college degree (48). ADNI participants were recruited at AD Research Centers, which are likely to attract people with concerns about memory and AD risk. There might, in turn, be a link between well-educated older adults with memory concerns and increased likelihood of progressing to Aβ-positivity. Further work will be necessary to fully explain this finding.

### Are the results driven by MCI cases?

We considered that the present results might be driven by the 47.3% of the sample diagnosed with MCI at baseline. However, ORs were in the direction of greater magnitude among cognitively normal when analyzed separately from MCI participants. It is also worth emphasizing that the results for the majority (52.7%) of the sample do not challenge the proposed AD continuum because these non-MCI individuals did not have cognitive impairment prior to reaching Aβ-positivity. Rather, differences that were present within the range of normal cognitive function were informative about who is more likely to become Aβ-positive.

### Implications for study participant selection

Use of Aβ-positivity as inclusion criteria should be context dependent. Defined cut-points are necessary for clinical diagnosis and in scenarios such as clinical trials targeting existing Aβ pathology. Including only dichotomously-defined, biomarker-confirmed MCI cases will reduce the number of false-positive diagnoses and provide more certainty that cognitive deficits arise from AD pathology. Our results suggest that early cognitive testing may also hold utility as a screening tool for identifying who should receive biomarker assessments to more directly assess disease etiology or suitability for clinical trials. However, it will exclude Aβ-negative MCI cases who may later enter the AD continuum upon progression to Aβ-positivity. If the goal of a study is to understand the earliest stages of the AD continuum, it will be important to capture individuals who demonstrate putative atypical disease progression to better detect and identify sources of variability.

### Summary

Despite much evidence for the standard model of biomarker and cognitive trajectories, the current results demonstrate the complex nature of disease progression. Differences in cognition that predict future progression to Aβ-positivity may be driven by subthreshold pathology, perhaps suggesting a need to reconsider current biomarker thresholds or to focus more on approaches that measure Aβ accumulation. Additionally, higher levels of tau are associated with increased risk of becoming Aβ-positive, thus elevated tau should considered when identifying those at risk for developing AD. A subset of individuals with MCI but normal Aβ levels may similarly end up on the AD pathway as indicated by later progression to Aβ-positivity. Importantly, the results strongly suggest that cognition should not be considered important only as a late-stage endpoint of AD. Rather, even when cognitive function is still within the normal range, it can provide a sensitive, low-cost, non-invasive predictor of risk, potentially before current thresholds for Aβ-positivity are reached.

## ACKNOWLEDGMENTS

This work was supported by National Institute on Aging R01 AG050595 (W.S.K., M.J.L., C.E.F.), R01 AG022381 (W.S.K.), R01 AG059329 (sub PI C.E.F), R01 AG056410 (M.S.P) and K08 AG047903 (M.S.P). Data collection and sharing for this project was funded by the Alzheimer’s Disease Neuroimaging Initiative (ADNI) (National Institutes of Health Grant U01 AG024904) and DOD ADNI (Department of Defense award number W81XWH-12-2-0012). ADNI is funded by the National Institute on Aging, the National Institute of Biomedical Imaging and Bioengineering, and through generous contributions from the following: AbbVie, Alzheimer’s Association; Alzheimer’s Drug Discovery Foundation; Araclon Biotech; BioClinica, Inc.; Biogen; Bristol-Myers Squibb Company; CereSpir, Inc.; Cogstate; Eisai Inc.; Elan Pharmaceuticals, Inc.; Eli Lilly and Company; EuroImmun; F. Hoffmann-La Roche Ltd and its affiliated company Genentech, Inc.; Fujirebio; GE Healthcare; IXICO Ltd.; Janssen Alzheimer Immunotherapy Research & Development, LLC.; Johnson & Johnson Pharmaceutical Research & Development LLC.; Lumosity; Lundbeck; Merck & Co., Inc.; Meso Scale Diagnostics, LLC.; NeuroRx Research; Neurotrack Technologies; Novartis Pharmaceuticals Corporation; Pfizer Inc.; Piramal Imaging; Servier; Takeda Pharmaceutical Company; and Transition Therapeutics. The Canadian Institutes of Health Research is providing funds to support ADNI clinical sites in Canada. Private sector contributions are facilitated by the Foundation for the National Institutes of Health (www.fnih.org). The grantee organization is the Northern California Institute for Research and Education, and the study is coordinated by the Alzheimer’s Therapeutic Research Institute at the University of Southern California. ADNI data are disseminated by the Laboratory for Neuro Imaging at the University of Southern California.

The funding agencies had no role in the design and conduct of the study; collection, management, analysis, and interpretation of the data; preparation, review, or approval of the manuscript; and decision to submit the manuscript for publication. Results of these analyses were reported at the 2019 Alzheimer’s Association International Conference and on the bioRxiv preprint server.

