## Supplemental for "Aβ-Positivity Predicts Cognitive Decline but Cognition Predicts Progression to Aβ-Positivity"

**Supplemental Material**

**Supplemental Methods**

*Rationale for logistic regression model versus generalized linear mixed-effects model*

Our question of interest was to identify individuals at elevated risk of entering the Alzheimer’s disease continuum as determined by progression to β-amyloid (Aβ) positivity. We therefore wished to consider all individuals who progress to Aβ-positivity equivalently, regardless of time to progression, as they are all at elevated risk and could therefore be appropriate candidates for intervention. We provide a hypothetical example to illustrate our motivation for choosing the logistic regression approach:

| Visit # | Participant 1 | Participant 2 | Participant 3 |
| --- | --- | --- | --- |
| 1 | 0 | 0 | 0 |
| 2 | 1 | 1 | 0 |
| 3 | 1 | NA | 0 |
| 4 | 1 | NA | 0 |
| 5 | 1 | NA | 1 |

The table above presents three hypothetical participants, and we will assume they all had the same low scores on baseline cognitive composites. A 0 indicates that they were Aβ-negative, and a 1 indicates Aβ-positive. NA indicates no data from that timepoint.

We chose to conduct a logistic regression with the outcome variable indicating whether an individual progressed from Aβ-negative at baseline to Aβ-positive at any point during follow-up or remained Aβ-negative throughout. Under the logistic regression model, all three of the above participants will be treated identically as they each will have the same value for their dependent variable (i.e., they are all classified as converters). Coefficients from this model can be interpreted as reflecting risk at the unit of the individual. In other words, is an individual at elevated risk, regardless of *when* progression occurs?

This approach was chosen rather than a generalized linear mixed-effects model (GLMM) with the logit, or a logistic regression for longitudinal data that includes an outcome variable indicating Aβ status at each timepoint. Although the logistic regression model includes length of follow-up as a covariate, a benefit of the longitudinal GLMM is that it directly incorporates all timepoints, which may better address differences in follow-up time such as variation in conversion trajectories (timing and even reversion from Aβ-positive at one follow-up visit to Aβ-negative during a later visit to be included).

Under the longitudinal model, each row will contribute to the association between baseline cognition and Aβ-positivity. Coefficients of the fixed-effects from the GLMM would indicate odds of being Aβ-positive *at each timepoint*, which, along with the random-effects, allows for differentiation of subjects with different conversion profiles. Therefore, the participants above will differentially affect the coefficient, i.e., the effect of baseline cognition. Participant 1 will increase the effect of baseline cognition more than Participant 2 due to having multiple Aβ-positive timepoints. Participant 3 will decrease the effect of baseline cognition relative to the other two because they have multiple Aβ-negative timepoints. In contrast, because our interest was in identifying all individuals at elevated risk, the number of follow-up assessments after conversion is not relevant and should not be used to alter this level of risk. The logistic regression model addresses this specific question with easy-to-interpret results.

*Rationale for survival analysis*

This still leaves open the issue of differential follow-up times between individuals that convert versus those who remain stable. It may be that if individuals who remained Aβ-negative were followed for longer, they would eventually become Aβ-positive. Therefore, we conducted survival analyses that more directly address this issue as well as timing of conversion by testing the association between baseline cognitive performance and time to event (either conversion to Aβ-positive or censored at last follow-up). In this case, Participants 1 and 2 from the example above will be considered equivalent. That is, they both convert at time 2, and the number of follow-ups beyond this event does not matter. Participants 3 is treated differently under this model in that they have a longer time to event and would weaken the effect of baseline cognition relative to the other two participants. From a conceptual standpoint, this model splits the difference between the other two models. It more directly addresses the issue of follow-up time and, consistent with our goals, the number of Aβ-positive assessments following conversion does not alter an individual’s risk.

**Supplemental Tables & Figures**

**Supplemental Table S1.** **Baseline sample characteristics of individuals with mild cognitive impairment with converted to Aβ-positivity versus remained Aβ-negative.** Mean (SD) presented for continuous variables, count (%) presented for categorical variables. An asterisk indicates a significant (p < 0.05) difference between the two groups.

|  | **MCI Aβ-stable** | **MCI Aβ-converter** |
| --- | --- | --- |
| n | 117 | 21 |
| Age | 70.46 (8.08) | 70.54 (8.31) |
| Gender, male | 63 (53.8%) | 10 (47.6%) |
| APOE-ε4+ status | 20 (17.1%) | 8 (38.1%) |
| Education* | 15.96 (2.56) | 17.43 (2.31) |
| Length of follow-up (years)* | 3.22 (1.33) | 3.44 (1.21) |
| Baseline CSF Aβ* | 1474.67 (245.65) | 1269.88 (278.38) |
| Baseline CSF P-tau* | 18.86 (7.71) | 23.28 (7.82) |

**Supplemental Figure S1.** **Baseline cognitive performance and risk of progression to Aβ-positivity.** Results of two Cox proportional hazard models using A) the ADNI Memory composite (ADNI_MEM) and B) the Preclinical Alzheimer Cognitive Composite (PACC). Measures are all taken from baseline and predict time to event (conversion to Aβ-positivity or censored at last follow-up). Cognitive scores were converted to z-scores and reverse coded such that higher scores indicate poorer performance. Hazard ratios are presented with asterisks indicating significant estimates (*p<0.05, **p<0.01, ***p<0.001). Lines represent 95% confidence intervals.

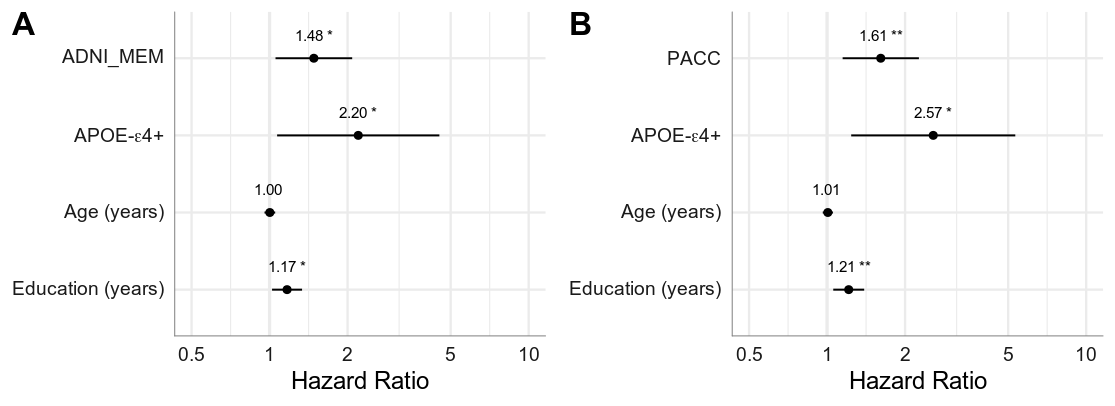

**Supplemental Figure S2.** **Baseline cognitive performance and continuous measures of CSF Aβ and p-tau associated with risk of progression to Aβ-positivity.** Results of two Cox proportional hazard models using A) the ADNI Memory composite (ADNI_MEM) and B) the Preclinical Alzheimer Cognitive Composite (PACC). Measures are all taken from baseline and predict time to event (conversion to Aβ-positivity or censored at last follow-up). Cognitive scores were converted to z-scores and reverse coded such that higher scores indicate poorer performance. CSF Aβ and P-tau were entered as continuous variables. Both measures were z-scored and CSF Aβ was reverse coded such that higher values on both indicates abnormality. Hazard ratios are presented with asterisks indicating significant estimates (*p<0.05, **p<0.01, ***p<0.001). Lines represent 95% confidence intervals.

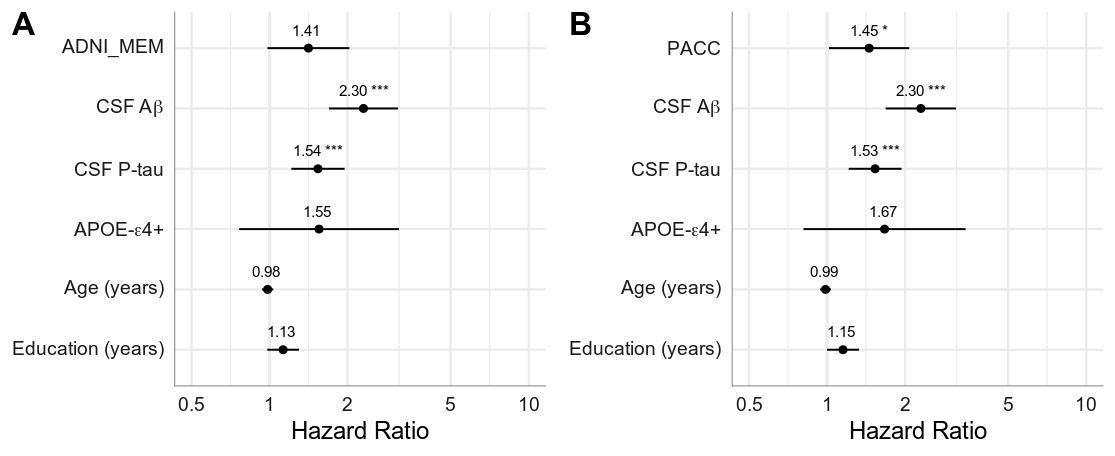
